# The contrasting response to drought and waterlogging is underpinned by divergent DNA methylation programs associated with gene expression in sesame

**DOI:** 10.1101/362905

**Authors:** Komivi Dossa, Marie Ali Mmadi, Rong Zhou, Qi Zhou, Mei Yang, Ndiaga Cisse, Diaga Diouf, Linhai Wang, Xiurong Zhang

**Affiliations:** Oil Crops Research Institute of the Chinese Academy of Agricultural Sciences, Key Laboratory of Biology and Genetic Improvement of Oil Crops, Ministry of Agriculture, No.2 Xudong 2nd Road, Wuhan 430062, China; Centre d’Etudes Régional Pour l’Amélioration de l’Adaptation à la Sécheresse (CERAAS), Route de Khombole, Thiès, BP 3320, Senegal; Laboratoire Campus de Biotechnologies Végétales, Département de Biologie Végétale, Faculté des Sciences et Techniques, Université Cheikh Anta Diop, BP 5005 Dakar-Fann, Code postal 10700, Dakar, Sénégal; College of Life Science, Hubei University, Wuhan, China

**Keywords:** sesame, drought, waterlogging, MSAP, DNA methylation pattern, gene expression regulation

## Abstract

DNA methylation is a heritable epigenetic mechanism that participates in gene regulation under abiotic stresses in plants. Sesame (*Sesamum indicum* L.) is typically considered a drought-tolerant crop but highly susceptible to waterlogging, a property attributed to its presumed origin in Africa or India. Understanding DNA methylation patterns in sesame under drought and waterlogging conditions can provide insights into the regulatory mechanisms underlying its contrasting responses to these principal abiotic stresses. Here, we combined Methylation-Sensitive Amplified Polymorphism and transcriptome analyses to profile cytosine methylation patterns, gene expression alteration, and their interplay in drought-tolerant and waterlogging-tolerant sesame genotypes under control, stress and recovery conditions. Our data showed that drought stress strongly induced *de novo* methylation (DNM) whereas most of the loci were demethylated (DM) during the recovery phase. In contrast, waterlogging decreased the level of methylation under stress but during the recovery phase, both DM and DNM were concomitantly deployed. In both stresses, the differentially expressed genes (DEGs) were highly correlated with the methylation patterns. We observed that DM was associated with the up-regulation of the DEGs while DNM was correlated with the down-regulation of the DEGs. In addition, we sequenced 44 differentially methylated regions of which 90% overlapped with the promoters and coding sequences of the DEGs. Altogether, we demonstrated that sesame has divergent epigenetic programs that respond to drought and waterlogging stresses. Our results also highlighted the possible interplay among DNA methylation and gene expression, which may modulate the contrasting responses to drought and waterlogging in sesame.

## Introduction

Sesame (*Sesamum indicum* L.) is a traditional oilseed crop widely grown in tropical areas (Bedigian, 2004). Over the past few years, growing attention has been paid to the crop because of the discovery of the health-promoting effects of its oil (Pathak et al., 2014). On the other hand, as a hardy crop able to survive in extreme climatic conditions, sesame production provides an opportunity to valorize marginal lands and represents an important source of income for small-scale farmers in developing countries. Globally, sesame production is increasing and the growing area is markedly expanding. Nonetheless, in the different growing regions, sesame has a very weak productivity and low yield, mainly due to the negative effects of abiotic stresses. Therefore, understanding the mechanisms of abiotic stress tolerance for improvement towards higher productivity and yield has become a hot topic in current sesame research (Dossa et al., 2017a). Two principal abiotic stresses including drought and waterlogging affect sesame productivity (Wang et al., 2016a). Drought stress is mainly significant in the arid and semi-arid areas of Africa, America and Asia, but, in south and East Asia, waterlogging represents the major threat for the sesame production. Sesame is typically considered a drought tolerant crop (Langham, 2007). However, intense and prolonged drought stress limits sesame plant growth, impairs flower production, reduces the formation of capsule and seed and ultimately, affects seed yield (Hassanzadeh et al., 2009; Sun et al., 2010; Boureima et al., 2012; Dossa et al., 2017b). When it occurs at the seedling stage, prolonged drought stress can result in increased plant mortality. Conversely, sesame is highly susceptible to waterlogging stress (Wang et al., 2012). While the crop can sustain drought stress for several days, few hours of flooding leads to catastrophic plant mortality and yield loss even in the case of waterlogging-tolerant varieties. The contrasting responses to drought and waterlogging suggest the existence of different underlying molecular mechanisms in sesame. It is speculated that being cultivated for millennia in drought-prone environments, sesame may have been genetically shaped with a sophisticated and heritable molecular mechanism for drought adaptation.

One of the molecular mechanisms worth investigating relates to environmentally induced epigenetic modifications. In fact, methylation of the DNA is the most stable and heritable epigenetic modifications which has been associated with regulation of gene expression and response to environmental stresses in plants (Paun et al., 2010; Dowen et al., 2012; Xia et al., 2017). It results in the covalent addition of a methyl group to the fifth position of the aromatic ring in cytosine. The cytosine base may be methylated by DNA methyltransferases (DNMTs) (Goll et al., 2003), or demethylated by REPRESSOR OF SILENCING 1 (ROS1), DEMETER (DME), and DEMETER like-proteins (DME 2,-3) (Zeng et al., 2008) in conjunction with environmental and developmental cues (Baulcombe and Dean, 2014). In plants, DNA methylation occurs commonly within three sequence contexts: CG, CHG and CHH (where H is A, C, or T); however, it varies depending on the level and pattern found within different genomic regions (Chwiaslkowska et al., 2017). Among these three cytosine contexts, CpG dinucleotides are typically clustered around the regulatory region of a gene, especially in the promoter and first exon, which can impact its transcriptional regulation (Garg et al., 2015).

Under stress, DNA methylation patterns depend on the plant species, the tissues and the specific type of stress. For example, total methylation increases under salt stress in alfafa (Al-Lawati et al., 2016) but decreases in salt-sensitive rapessed (Marconi et al., 2013). It has been reported that drought stress induces both demethylation and *de novo* methylation of DNA throughout the genome of barley (Chwialkowska et al., 2016) and ryegrass (Tang et al., 2014). Under drought treatment, a decrease of DNA methylation was observed in leaf tissues of two faba bean genotypes (Abid et al., 2017). Moreover, Bednarek et al. (2017) observed an elevated demethylation in both non-tolerant and tolerant plants, with *de novo* methylation occurring less frequently than demethylation under Aluminum stress in triticale lines.

The methylation-sensitive amplified polymorphism (MSAP) technique developed by Reyna-López et al. (1997) is an adaptation of amplified fragment length polymorphism analysis and has proven to be a powerful tool for analyzing DNA methylation. The MSAP technique has been applied to study CpG methylation in the genome of plants, somaclonal epigenetic variation, cytosine methylation during various developmental stages and resistance to biotic and abiotic stresses (Ashikawa, 2001; Matthes et al., 2001; Portis et al., 2004; Sha et al., 2005, Ruiz-Garcia et al., 2005; Tang et al., 2014; Zhang et al., 2015; Bednarek et al., 2017; Abid et al., 2017). Different approaches have been proposed to interpret MSAP outputs (Schulz et al., 2013; Fulneček and Kovařik, 2014), but none takes into account the multiple events reflecting various methylation patterns that must take place simultaneously to explain the differences in individual digestion patterns between control and stressed materials. Recently, Bednarek et al. (2017) introduced an efficient theoretical model for the quantification of cytosine methylation patterns at restriction sites of the isoschizomers *Hpa*II and *Msp*I which evaluates demethylation, *de novo* methylation, and preservation of methylation status between control and stressed samples.

While the contribution of DNA methylation to plant performance under abiotic stress has been well studied in other major crops, no study has been done in sesame. Previously, our group sequenced the whole transcriptome under drought (Dossa et al., 2017c,d) and waterlogging (Wang et al., 2016a) at different time points. These data represent an important resource to get insight into the interplay among DNA methylation and gene expression regulation in sesame. The main objective of the present investigation is to test the hypothesis that drought and waterlogging stress induce divergent DNA methylation patterns as the basis of the contrasting responses of sesame.

## 2. Materials and Methods

### 2.1. Plant materials and stress treatment

Two genotypes of sesame (*Sesamum indicum* L.) were obtained from the China National Genebank, Oil Crops Research Institute, Chinese Academy of Agricultural Sciences and used in this experiment. The genotype ZZM0635 displays a tolerance to drought stress (Dossa et al., 2017c,d) while the genotype Zhongzhi No.13 is relatively waterlogging tolerant (Wang et al., 2016a). The seeds were sown in pots (25 cm across and 30 cm deep) containing 6 kg of loam soil mixed with 10% vermiculite. The experiment was conducted under shelter in natural conditions with the mean temperature of 31/27°C day/night. A completely randomized blocking design with 4 replicates was employed and plants were watered normally under the optimum soil volumetric water content (vwc) of 35%. The soil moisture was measured manually in each pot using a Moisture Meter *Takeme* over the entire experiment. At the flowering stage, the irrigation was suspended for 11 days (DS) in 1/3 of the pots with the soil volumetric water content falling from 35% to 6%. The plants displayed heavy wilting signs, thereafter, they were allowed to recover for 4 days (DR) by re-supplying irrigation to reach the optimum soil volumetric water content (35%) according to the experimental descriptions of Dossa et al. (2017d) (Fig. 1). For the waterlogging application, 1/3 of the pots were flooded by standing in a plastic bucket filled with tap water to 3 cm above the soil surface and maintained for 9 h (WS) according to the experimental descriptions of Wang et al. (2016a). Under stress, plants showed moderate wilting signs as presented in Fig. 1. Then, water was drained from the pots to allow the plants to recover for 20 h (WR). Meanwhile, the control plants (D-CK and W-CK) were kept under normal irrigation condition (35% vwc) throughout the whole experiment. The roots of 3 stressed plants were harvested individually under stress application and during recovery and those of the control plants were sampled at the same periods.

**Fig. 1.**
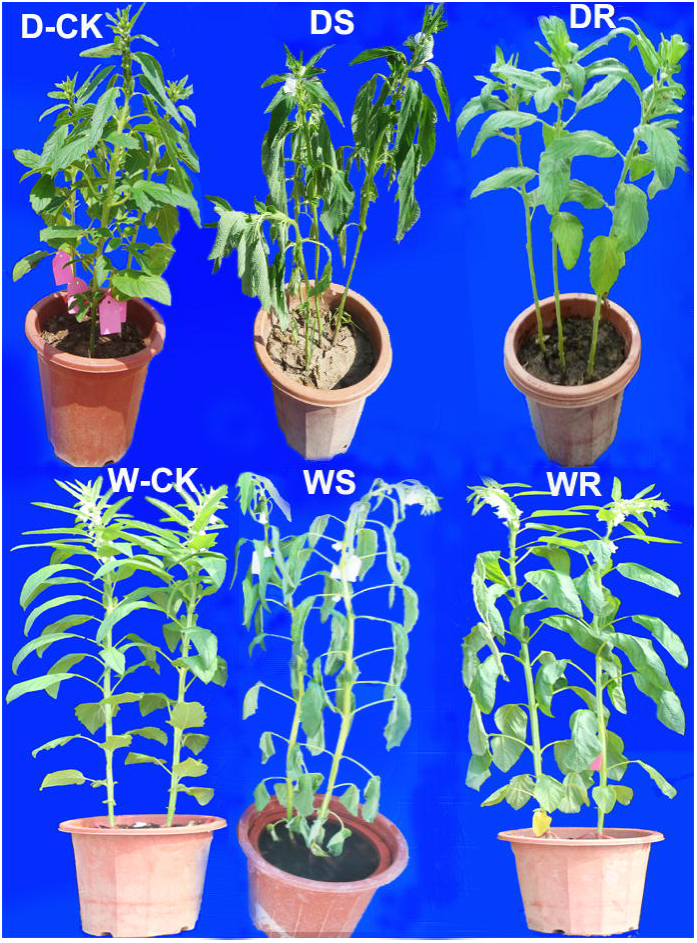
Phenotype characteristics of drought tolerant (D) and waterlogging tolerant (W) sesame genotypes under control (CK) stress (S) and recovery (R) treatments.

### 2.2. DNA extraction and MSAP epigenotyping

DNA was extracted from the root samples using the hexadecyltrimethylammonium bromide CTAB method (Dossa et al., 2016). The quality of the DNAs was checked on 1% agarose gel and the quantity was evaluated on the ultraviolet spectrophotometer. Aliquots were diluted to the final concentration of 300 ng.µl^−1^. The extracted DNAs were subjected to a Methylation-Sensitive Amplified Polymorphism (MSAP) analysis following the method of Xiong et al. (1999) with slight modifications. The technique is based on the use of the isoschizomers *Hpa*II and *Msp*I that differ in their sensitivity to methylation of their recognition sequences. Both enzymes recognize the tetranucleotide sequence 5’-CCGG-3’, but their action is affected by the methylation state of the external or internal cytosine residues. 2 µL of each DNA sample (300 ng) was digested with 0.5 µL *EcoR*I-10 U with 13.5 µL deionized H_2_O and 4 µL 10× Tango buffer (Thermo, USA) at 37°C for 2 h, before deactivation by heating at 65°C for 20 min. Then, the digested DNA fragments were subjected to *Hpa*II-10 U and M*sp*I-10 U digestion into two separated series at 37°C overnight. The restriction enzymes were deactivated by heating at 80°C for 15 min. The ligation was performed in a final volume of 10 µl including 5 µl enzyme-digested products, 1 µL of 5 pmol *EcoR*I adapter (Table S1), 1 µL of 50 pmol *Hpa*II/*Msp*I adapter (Table S1), 3 µL deionized H_2_O, 1 µL 10× T4 ligase buffer (Promega, USA) and 0.5 µL T4 DNA ligase (5U.µl^−1^) and 1 µL ATP incubated at 37°C overnight.

Pre-selective amplification was performed in a 20 µL reaction volume with 0.5 µl *EcoR*I primer (10 mM) and 0.5 µl *Msp*I/*Hpa*II primer (10 mM), 2.5 µl restriction-ligation DNA, 8 µl of 2× Reaction Mix (Tiangen Biotech, Beijing, China) supplied together with the dNTPs and MgCl_2_, 8 µl deionized H_2_O and 0.5 µl of 1 U Taq polymerase. The pre-selective amplification was conducted with the following temperature cycling conditions: 1 cycle at 94°C for 5 min; 30 cycles at 94°C for 30 s, 56°C for 30 s, and 72°C for 15 s, and finally one cycle at 72°C for 7 min prior to 30 min incubation at 60°C.

Selective amplification was conducted in a 20 µL volume including 3 µL of 10-fold diluted pre-amplified PCR products, 8 µL of 2× Reaction Mix (Tiangen Biotech, Beijing, China) supplied together with the dNTPs and MgCl_2_, 0.5 µl of primers (10 mM), 7 µl deionized H_2_O and 1 µl of 1 U Taq polymerase. A total of 60 primer combinations were tested on 4 DNA samples and 25 primer combinations (E_*n*_/HM_*n*_) displaying clear PCR profiles were finally retained for MSAP epigenotyping (Table S1). The PCR amplification reactions were performed using touch-down cycles under the following conditions: 94°C for 5 min; 13 touch-down cycles of 94°C for 30 s, 65°C (subsequently reduced each cycle by 0.7°C) for 30 s and 72°C for 30 s; 23 continued cycles of 94°C for 30 s, 56°C for 30 s and 72°C for 15 s, and finally extension at 72°C for 7 min. The PCR products of selective amplifications were separated using 6% polyacrylamide gel electrophoresis at 50 W for 1 h 30 min. Gels were silver stained using the method described by Benbouza et al. (2006). MSAP analysis was performed twice for each primer combination.

### 2.3. Data scoring and analysis of MSAP profiles

The MSAP profiles showing reproducible results between replicates were transformed into a binary matrix, with 1 as a presence of band and 0 the absence of band from either *EcoR*I/*Msp*I or *EcoR*I/*Hpa*II digestion and analyzed according to the protocol developed by Bednarek et al. (2017). In general, these fragments could be divided into four types representing four types of DNA methylation status of the restriction sites (5′-CCGG-3′): unmethylated (Type I, presence of the band in both enzyme combinations), hemi-methylated at the outer cytosine in one DNA strand (Type II, presence of the band only in digestion with *EcoR*I/*Hpa*II), fully-methylated at the internal cytosine in both DNA strands (Type III, presence of the band only in digestion with *EcoR*I/*Msp*I), and hyper-methylated with outer methylation at both DNA strands (Type IV, absence of band in both enzyme combinations) (Fulneček and Kovařík, 2014). Only 100-bp or longer PCR products were considered for analysis. Also, two main treatment comparisons were done in this study: Stress vs Control and Recovery vs Stress so as to understand epigenetic changes during stress and recovery, respectively. The resultant code of the binary matrix, expressed as 4 binary digits, describes the presence/absence of each fragment in the *EcoR*I*/Hpa*II and *EcoR*I*/Msp*I digests of DNA from 2 compared treatments. Theoretically, 16 permutations are possible and could be classified into demethylation (DM), *de novo* methylation (DNM), preservation of methylated sites (MSP), and preservation of non-methylated sites (NMSP) which are the 4 events that were investigated simultaneously in this study. We assume that these events are equally probable, then, the multiplication of the number of individual events participating in the explanation of a given 4-digit code (Table S2) by the number of MSAP profiles depicted by that code, followed by summation of events of the same kind and normalization of the data (expressed as percentages), correspond to the relative quantitative characteristics of these 4 events (Bednarek et al., 2017).

### 2.4. Sequencing of polymorphic MSAP fragments

The polymorphic MSAP fragments are relative to demethylation (DM) and *de novo* methylation (DNM). A total of 80 polymorphic bands (42 drought and 38 waterlogging MSAP bands) were excised from fresh gel and transferred into 0.6 mL tube. The excised gels were crushed and dissolved in 50 µL deionized H_2_O prior incubation at 50°C overnight. These bands were re-amplified with the appropriate selective primer combinations. Sizes of the PCR products were checked by agarose gel electrophoresis and 57 positive PCR products were sent for sequencing at Tsingke (www.tsingke.net). Homology analysis with the reference sesame genome sequence was performed via the BLASTn search program with a cut off E-value ≤1×10^−40^ on Sinbase (http://ocri-genomics.org/Sinbase/index.html) (Wang et al., 2014).

### 2.5. RNA extraction and qRT-PCR analysis

Total RNAs from root samples were extracted with the Easy Spin RNA kit (Aidlab, Beijing, China) following descriptions by Dossa et al. (2017d). The quantity and quality of RNA samples were assessed by 1% agarose gel electrophoresis and on the ultraviolet spectrophotometer measurement of the A260/A280 ratio. For cDNA synthesis, 1.5 µg of RNA was reverse transcribed using the Superscript III reverse transcription kit (Invitrogen, Carlsbad, CA, USA), according to the manufacturer’s instructions. We designed primers to amplify the differentially methylated genes using the Primer Premier 5.0 software (Lalitha, 2000) (Table S3). The qRT-PCR was conducted on a Roche Lightcyler® 480 instrument using the SYBR Green Master Mix (Vazyme), according to the manufacturer’s protocol. Each reaction was performed using a 20 µL mixture containing 10 µL of 2× ChamQ SYBR qPCR Master Mix, 6 µL of nuclease-free water, 1 µL of each primer (10 mM), and 2 µL of 4-fold diluted cDNA. All of the reactions were run in 96-well plates and each cDNA was analyzed in triplicate. The following cycling profile was used: 95°C for 30 s, followed by 40 cycles of 95°C/10 s, 60°C/30 s. Each reaction was conducted in biological triplicates and the sesame *Histone H3.3* gene (*SIN_1004293*) was used as the endogenous control gene (You et al., 2018). The method developed by Livak and Schmittgen (2001) was employed for the data analysis.

### 2.6. RNA-seq data analysis

Raw data of drought and waterlogging RNAseq experiments were retrieved from GeneBank Short Read Archive (SRA) with the accession numbers SAMN06130606 and SRR2886790, respectively. The sequencing reads containing low-quality were cleaned with FastQC (http://www.bioinformatics.babraham.ac.uk/projects/fastqc/). The clean reads were mapped to the sesame reference genome v2.0 (Wang et al., 2016b) using HISAT2 (Kim et al., 2015); the program StringTie (Pertea et al., 2016) was used for transcript assembly. The Cufflinks 2.0 software (Trapnell et al., 2010) was used to calculate the gene expression level for each sample expressed as fragments per kilobase of transcript per million fragments mapped (FPKM). The differentially expressed genes (DEG) were detected as described by Tarazona et al. (2011) based on the parameters: Fold change > = 2.00 and Probability > = 0.8. The DEGs were identified by comparing: DSvsCK, DRvsDS for drought stress and WSvsCK, WRvsWS for waterlogging stress. The log_2_ transformed FPKM values were used to construct heatmap using the MEV software (Saeed et al., 2006).

## 3. Results

### 3.1. Global DNA methylation levels under control, drought, waterlogging and recovery treatments in sesame

Cytosine methylation patterns in the root of drought-tolerant and waterlogging sesame genotypes under control (CK), stress (DS, WS) and recovery (DR, WR) conditions were assessed using 25 primer combinations (Table S1). Similar numbers of clear and reproducible bands were successfully revealed in drought (559) and waterlogging (533) conditions. Furthermore, most of the CCGG sites were shown to be largely methylated with the values ranging between 63.86% and 48.59% (Table 1). In the control treatment, we observed slight variations in the numbers of methylated sites between drought and waterlogging. However, in the stress treatment, drought strikingly increased the methylation level while it was decreased under waterlogging stress when levels of DNA methylation were compared with those in the respective unstressed control. At the recovery stage, the levels of methylation tended to reach those observed under control conditions. Further analyses showed that fully-methylated bands were more predominant that the hemi-methylated ones in all conditions, except for WR.

**Table 1.** Global DNA methylation patterns in root of sesame genotypes under control, stress (drought and waterlogging) and recovery treatments

| MSAP band type | CK | DS | DR | CK | WS | WR |
| --- | --- | --- | --- | --- | --- | --- |
| I | 266 | 202 | 278 | 248 | 274 | 258 |
| II | 105 | 68 | 66 | 83 | 64 | 165 |
| III | 172 | 47 | 92 | 140 | 181 | 60 |
| IV | 16 | 242 | 123 | 62 | 14 | 50 |
| Total amplified bands | 559 | 559 | 559 | 533 | 533 | 533 |
| Total methylated bands | 293 | 357 | 281 | 285 | 259 | 275 |
| MSAP (%) | 52.42 | 63.86 | 50.27 | 53.47 | 48.59 | 51.59 |
| Fully-methylated (%) | 33.63 | 51.70 | 38.46 | 37.90 | 36.59 | 20.64 |
| Hemi-methylated (%) | 18.79 | 12.16 | 11.81 | 15.57 | 12.00 | 30.95 |

Type II = hemi methylated bands and types III + IV are full methylated bands. Total methylated bands = II + III + IV.

### 3.2. DNA methylation alterations under stress and recovery treatments

To evaluate the impact of stress and recovery treatments on DNA methylation in sesame, the differentially methylated DNA bands between DSvsCK, DSvsDR, WSvsCK and WRvsWS were classified into 4 events including demethylation (DM), *de novo* methylation (DNM), preservation of methylated sites (MSP), and preservation of non-methylated sites (NMSP) according to the method developed by Bednarek et al. (2017). A total of fifteen 4-bit codes representing the pattern changes at a given cytosine between compared treatments, were identified in this study (Table 2). In fact, the code ‘‘0000’’ was not taken into account because this pattern cannot be easily recognized. Interestingly, the most frequent codes in drought stress compared with the control condition were 0100, 1000 which were totally different from the predominant pattern changes from drought to recovery phase (0001 and 0010). Conversely, the most induced methylation alteration under waterlogging stress and the recovery phase are the cases corresponding to the codes 0100 and 0001 (Table 2). Table 3 presents the converted codes into events of the corresponding type (based on Table S2), and their relative quantification. The results indicated that the imposition of drought stress induced principally DNM (41.97%) and to a lesser extent DM (22.03%) in the sesame genotype. In addition, DNM at the CHG sites was the most frequent (30.92%) while DM affected predominantly the CG sites (13.65%). About 36% of loci were not affected by drought stress at the epigenetic level. From DS to DR, an opposite trend was observed. Most of loci were demethylated with the majority harboring a CHG site. Moreover, few sites underwent DNM and about 37% of loci remained unchanged. These results suggest a divergent and drastic reprogramming of the methylation pattern from drought stress to the recovery stage in sesame. In the case of waterlogging, DM occurred at a higher proportion than DNM under stress. However, at the recovery phase, both DNM and DM were proportionally activated as epigenetic responses. In the two treatments, DNM-CHG and DM-CHG were the most represented types of methylation and demethylation, respectively (Table 3). Similarly as in drought conditions, 37% of loci were not affected by waterlogging and recovery treatments.

**Table 2.**
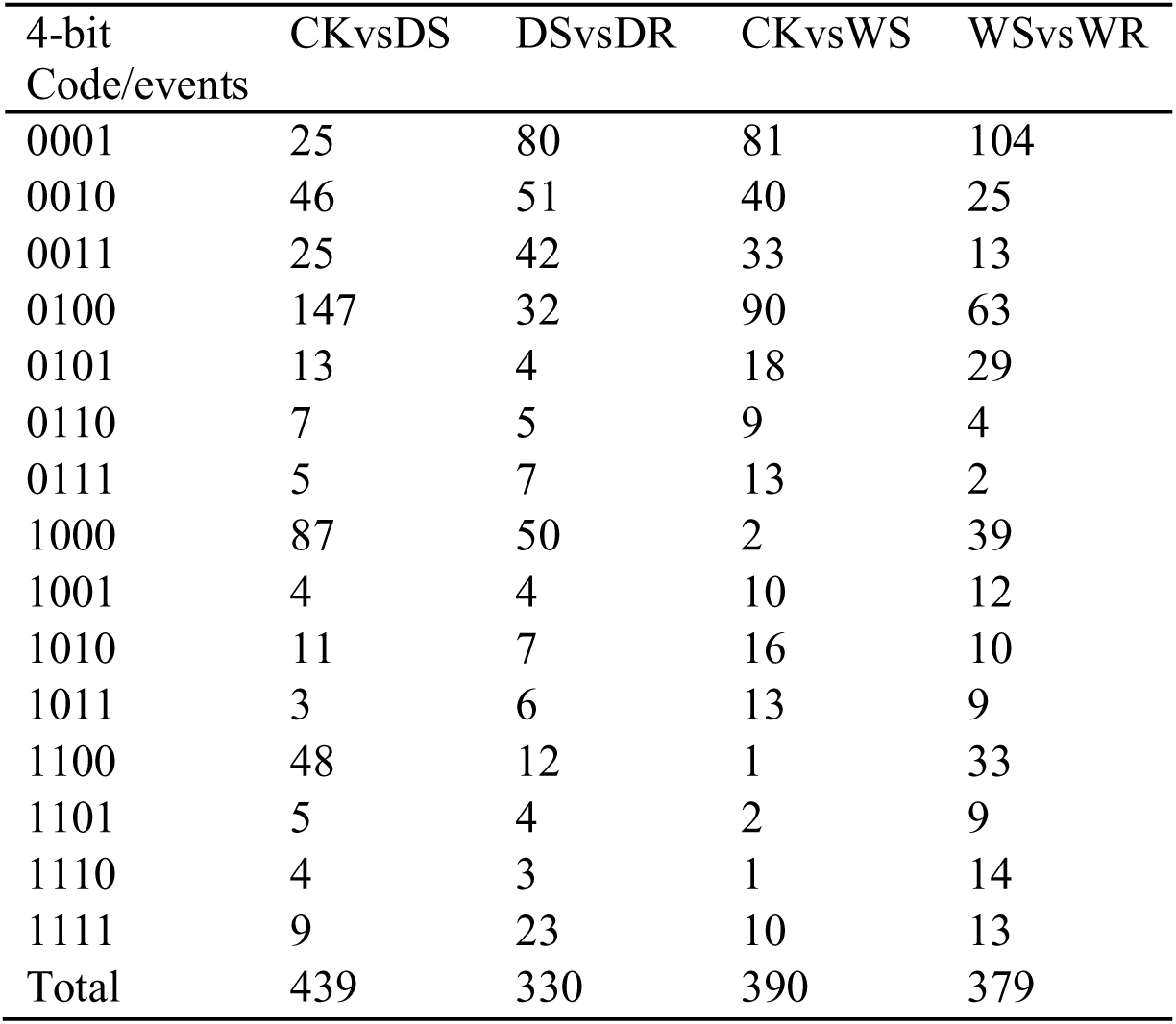
Profiles reflecting given MSAP 4-bit binary code evaluated among compared treatments

**Table 3.** Summary of the relative quantification of the DNA methylation patterns evaluated based on MSAP data

| Cases (events) | DSvsCK | DRvsDS | WSvsCK | WRvsWS |
| --- | --- | --- | --- | --- |
| DM | 1060 (22.03) | 1326 (39.29) | 1474 (35.00) | 1383 (31.40) |
| DNM | 2020 (41.97) | 804 (23.82) | 1140 (27.07) | 1336 (30.32) |
| MSP | 858 (17.83) | 565 (16.74) | 753 (17.88) | 784 (17.80) |
| NMSP | 874 (18.17) | 680 (20.15) | 845 (20.05) | 902 (20.48) |
| DNM-CHG | 1488 (30.92) | 380 (11.26) | 747 (17.74) | 725 (16.46) |
| DNM-CG | 532 (11.06) | 424 (12.56) | 393 (9.33) | 611 (13.87) |
| DM-CHG | 403 (8.37) | 924 (27.38) | 893 (21.20) | 967 (21.95) |
| DM-CG | 657 (13.65) | 402 (11.91) | 581 (13.79) | 416 (9.44) |
| Total | 4812 | 3375 | 4212 | 4405 |

Data in bracket are percentage

### 3.3. Comparative analysis of DNA methylation patterns and gene expression regulation under stress and recovery treatments

The relationship between DNA methylation variation and the gene expression regulation under drought and waterlogging stresses in sesame was assessed. Under drought stress, the analysis of the differentially expressed genes (DEGs) showed that 77% of DEGs were down-regulated whereas at the recovery stage, more than 80% of DEGs were up-regulated (Fig. 2a, b). These patterns of gene expression regulation correlate well with the DNA methylation changes (Fig. 2c). It implies that the high DNM observed under drought stress may induce a down-regulation of a large number of drought-responsive genes while the high DM observed at the recovery stage allowed for the resumption of drought-responsive gene expression. An inverse trend was observed in the waterlogging treatments with 60% of DEGs up-regulated under stress while, comparable numbers of DEGs were down- and up- regulated during the recovery stage (Fig. 2d,e). A similar reasoning about the correlation of the observed DNA methylation patterns and the gene expression alteration under waterlogging suggests that a high DM under stress was favorable for the up-regulation of waterlogging-responsive genes in the sesame genotype. We further scrutinized the common DEGs between DSvsCK and DRvsDS and found that 79% of down-regulated genes under stress were up-regulated during the recovery from drought damage while only 20% of DEGs experienced the opposite scenario (Fig. 3a). By analyzing the common DEGs between WSvsCK and WRvsWS, we observed that 37% of the down-regulated common DEGs under stress were up-regulated during the recovery stage while 58% of the common DEGs exhibited the opposite regulation pattern (Fig. 3b).

**Fig. 2.**
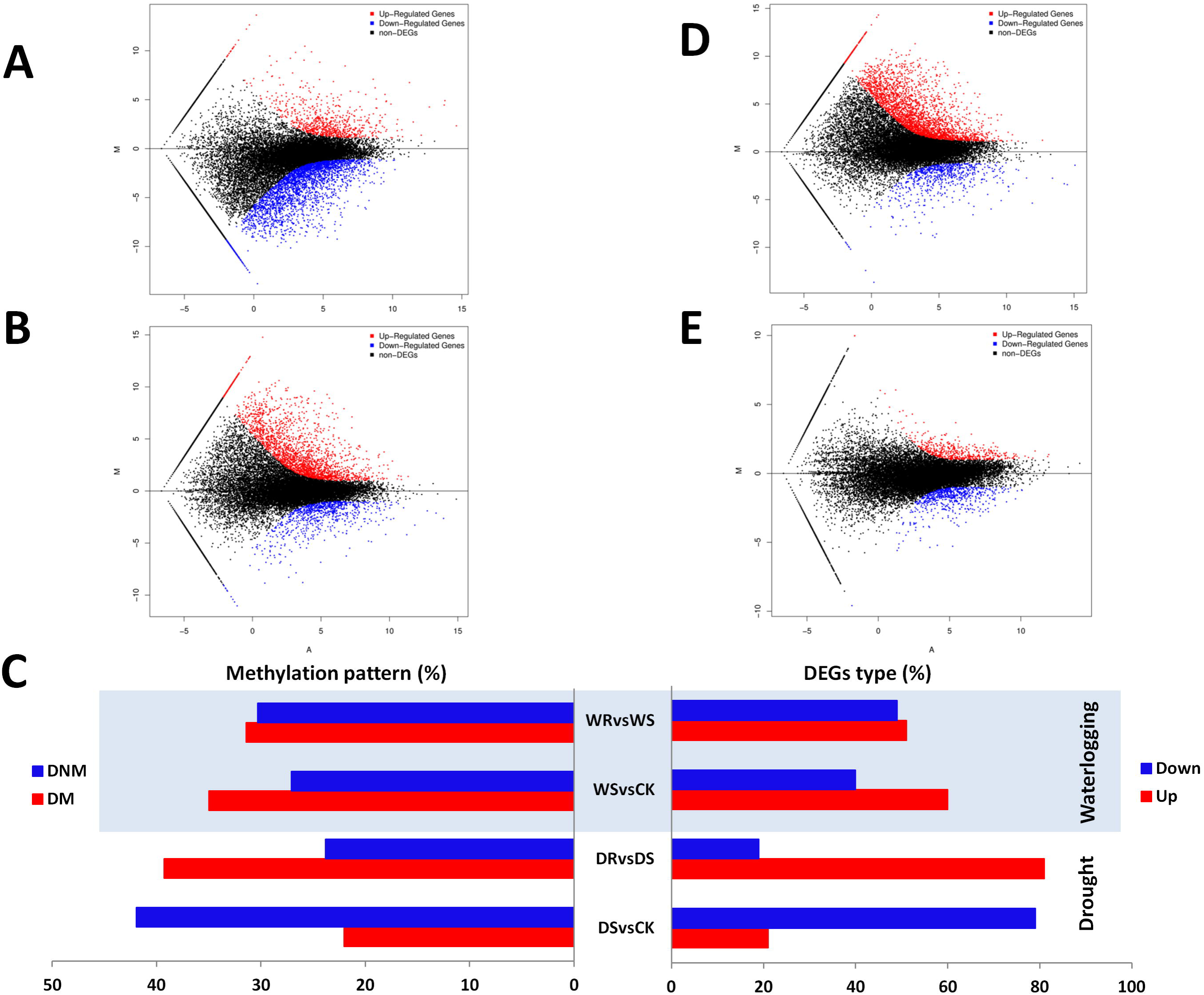
Gene regulation under drought and waterlogging and its correlation with DNA methylation. A. MA plot showing the differentially expressed genes (DEG) between drought stress (DS) and the control treatments (CK), B. MA plot showing the DEGs between drought recovery (DR) and drought stress treatments (DS), C. Positive correlation between DNA methylation patterns (*de novo* methylation (DNM) and demethylation (DM)) with gene expression regulation (up-regulation and down-regulation) in waterlogging and drought conditions, D. MA plot showing the DEGs between waterlogging stress (WS) and the control treatments (CK), E. MA plot showing the DEGs between waterlogging recovery (WR) and waterlogging stress treatments (WS).

**Fig. 3.**
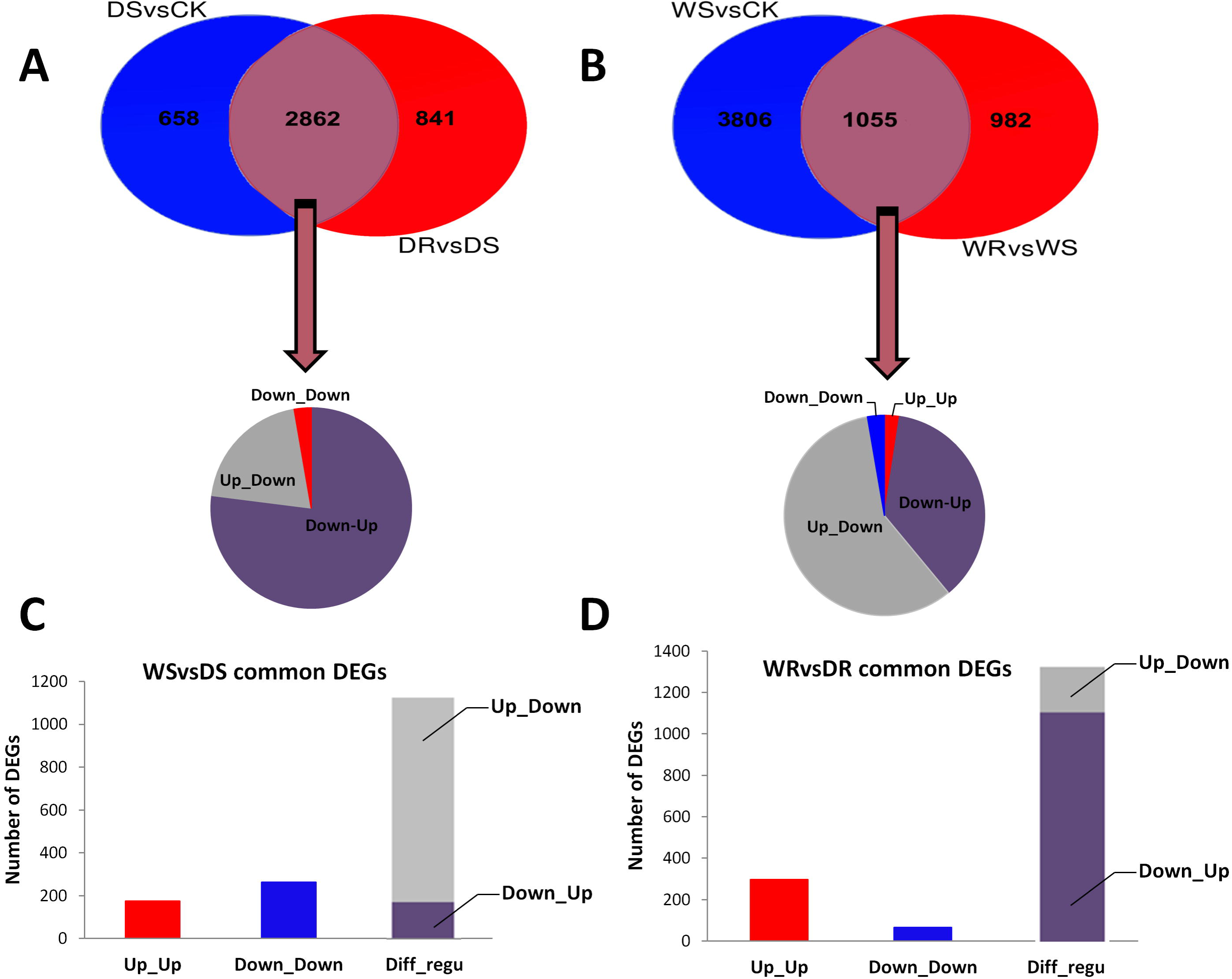
Regulation patterns of the common differentially expressed genes (DEG) between treatments and stresses. A. Shared and unique DEGs between drought stress (DSvsCK) and recovery (DRvsDS) and change in regulation status of the shared DEGs. Down_Down relates to DEGs constitutively down-regulated under drought stress and at the recovery phase; Up_Down relates to DEGs which were up-regulated under drought stress and down-regulated during the recovery phase; Down_Up relates to DEGs which were down-regulated under drought stress and up-regulated during the recovery phase. C. Comparison of DEGs between waterlogging and drought under stress (WSvsDS). Down_Down relates to DEGs constitutively down-regulated under drought stress and waterlogging stress; Up_Up relates to DEGs constitutively up-regulated under drought stress and waterlogging stress; diff_regu relates to DEGs that displayed contrasting regulation status between drought and waterlogging under stress. D. Comparison of DEGs between waterlogging and drought at the recovery stage (WRvsDR). Down_Down relates to DEGs constitutively down-regulated under drought recovery and waterlogging recovery; Up_Up relates to DEGs constitutively up-regulated under drought recovery and waterlogging recovery; diff_regu relates to DEGs that displayed contrasting regulation status between drought and waterlogging during recovery.

Moreover, we compared the DEGs between waterlogging and drought under stress (WSvsDS) and at the recovery (WRvsDR) stages. We identified 1526 and 1739 common DEGs for WSvsDS and WRvsDR, respectively. Within the common DEGs for WSvsDS, the majority (1087) displayed contrasting regulation patterns with most of these genes up-regulated in waterlogging stress while they were down-regulated under drought stress (Fig. 3c). In regard to the common DEGs at the recovery phases (WRvsDR), the majority also was conversely regulated between drought and waterlogging. In this case, however, most of these genes were down-regulated during the waterlogging recovery whereas they were found up-regulated during the drought recovery (Fig. 3d). Overall, these results indicated that drought and waterlogging induced divergent gene regulation in sesame strongly correlated with the DNA methylation patterns.

### 3.4. Analysis of polymorphic MSAP fragments and identification of differentially methylated genes

Out of the 57 sequenced polymorphic MSAP fragments, 44 bands including 24 for drought and 20 for waterlogging showed high similarities (97-100%) with the sesame genomic regions. The size of the excised bands ranged from 115 to 600 bp. The sequence analysis indicated that these successfully sequenced fragment termini have the CCGG site. Additionally, 17 fragments have one or more internal CCGG sites suggesting that the relative total methylation levels in sesame may be underestimated by the MSAP technique. Table 4 presents the methylation events (4-digit codes) of all the sequenced fragments and their associated genes. Interestingly, 40 differentially methylated genes were enlisted within the DEGs from drought and waterlogging treatments. In addition, 5 fragments overlapped with gene coding sequences (CDS) while the remaining fragments were located in the promoter region (UTR_5) of the associated genes. Functional annotation of the differentially methylated genes demonstrated that various classes of genes are methylated in response to drought and waterlogging stresses in sesame. The expression patterns of these differentially methylated genes were investigated using the transcriptome data in control, stress and recovery conditions (Fig. 4). The gene expression levels of all the differentially methylated genes changed from one treatment to another. In drought condition, the expression levels of most of the genes decreased from the control to the stress treatment, subsequently increasing from the stress to the recovery treatment (Fig. 4a). Meanwhile, in waterlogging condition, the expression level of most the differentially methylated genes were mainly induced during stress and at the recovery stage (Fig. 4b). Importantly, for drought and waterlogging differentially methylated genes, the patterns of gene expression changes matched well the methylation events represented by the 4-digit codes in Table 4. It is obvious that most of genes that experienced DNM were down-regulated while DM principally leads to the up-regulation of the gene expression level. We selected 16 and 14 differentially methylated genes from drought and waterlogging conditions, respectively, to further validate their expression changes using qRT-PCR. As shown in Fig. 4c,d, the qRT-PCR results corroborated well the transcriptome quantification of the expression level of the differentially methylated genes. These results demonstrated that the MSAP technique is an efficient approach to isolate stress-responsive genes. Altogether, our results highlighted an intimate correlation between DNA methylation pattern and gene expression regulation under drought and waterlogging in sesame.

**Fig. 4.**
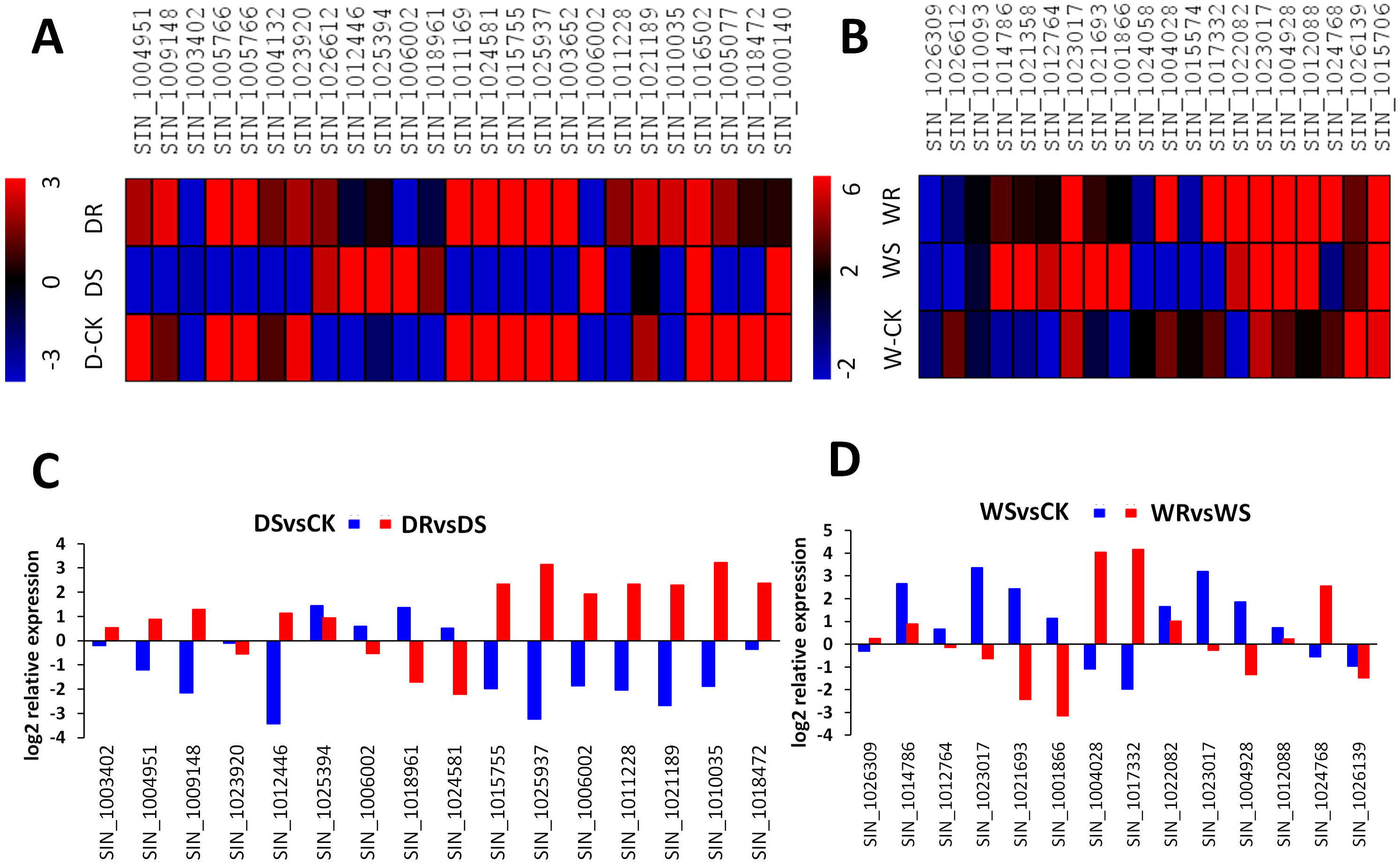
Expression analysis of the differentially methylated genes. A. a heatmap displaying the expression profile of 24 differentially methylated genes under control (D-CK), drought stress (DS) and drought recovery (DR) based on RNAseq data. B. a heatmap displaying the expression profile of 20 differentially methylated genes under control (W-CK), waterlogging stress (WS) and waterlogging recovery (WR) based on RNAseq data. The blue color depicts the weakly expressed genes while the red color depicts the highly expressed genes. C,D. qRTPCR validation of 16 and 14 selected gene expression under drought and waterlogging conditions, respectively. The blue bars correspond to the relative expression of the gene under stress compared with the control whereas the red bars correspond to the relative expression of the gene during the recovery phase compared with the stress.

**Table 4.**
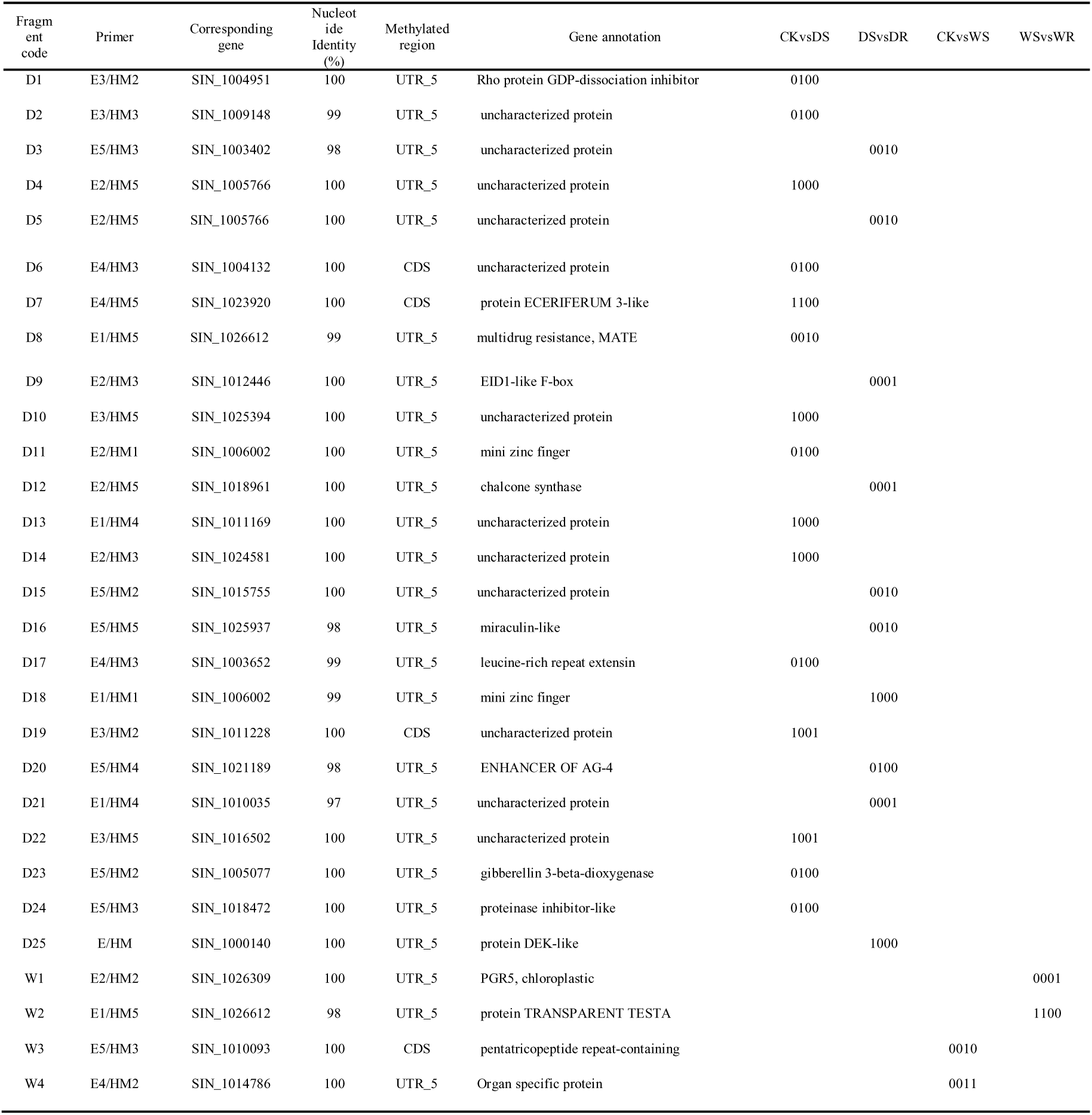
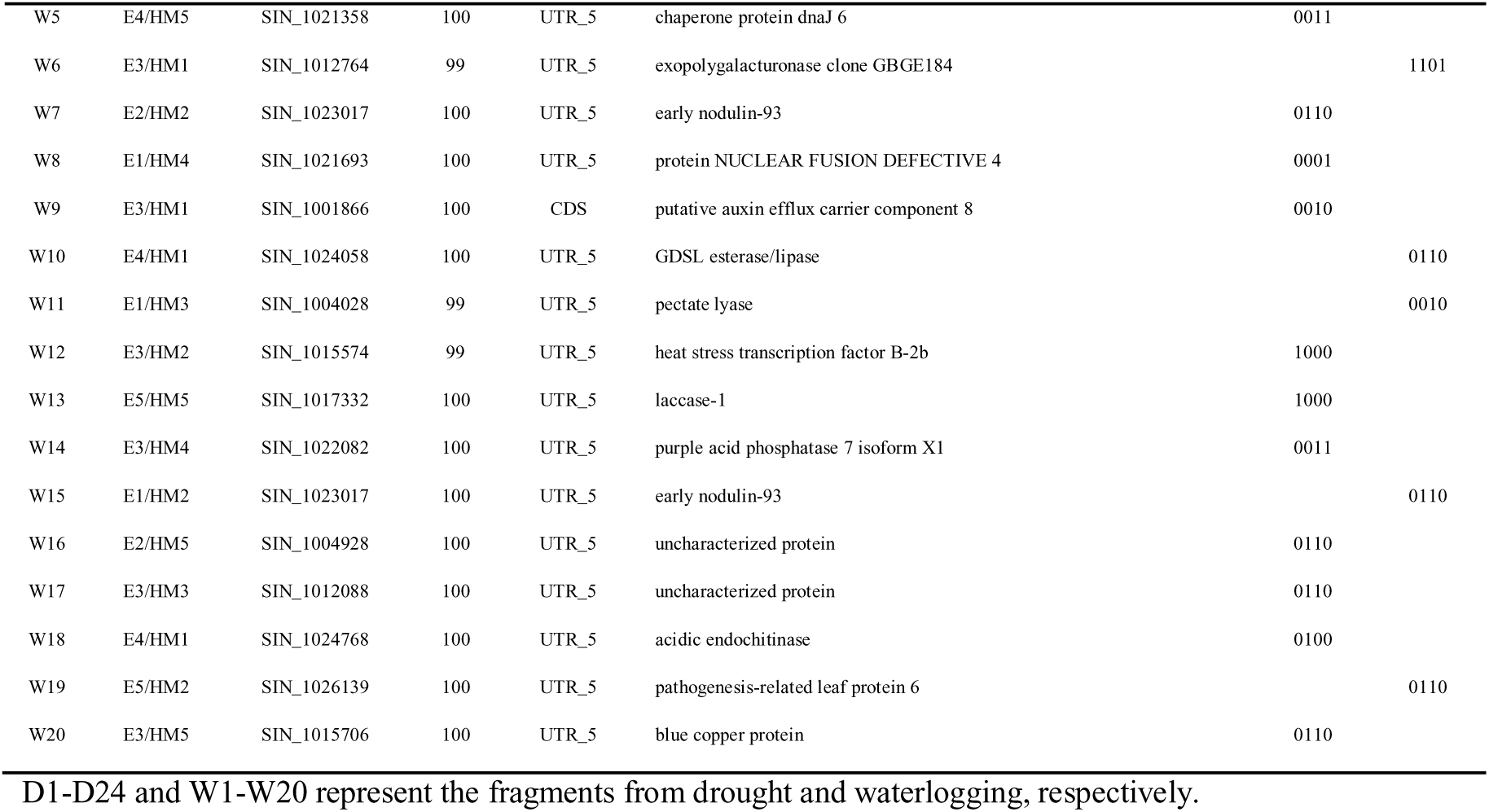
Blast results and methylation patterns of randomly selected polymorphic MSAP fragments

## 4. Discussion

As sessile organisms, plants respond to abiotic stresses by adjusting their physiological and developmental machinery through differentially regulated gene expression (Farooq et al., 2009). Mechanisms such as DNA methylation and demethylation of cytosine have been demonstrated to play a key role in this adjustment (Banerjee and Roychoudhury, 2017). In the present study, cytosine methylation analysis of sesame (~ 350 Mb) using the MSAP approach revealed that the level of methylation in the normal growth condition of two different genotypes was similar (52-53%). This level of methylation is close to those of ryegrass (57%), rapessed (46%), faba bean (41%) but obviously higher than those of rice (31%), *Arabidopsis thaliana* (30%), maize (35%) (Wang et al., 2015; Dai et al., 2015; Cokus et al., 2008; Karan et al., 2012; Shan et al., 2013; Tang et al., 2014). Using the MSAP sequencing technique, Pan et al. (2012) reported that the overall level of DNA methylation was about 70% in *Triticum aestivum*. Hence, these observations suggested that DNA methylation is a direct function of the plant species, particularly, the genome size and structure.

Under stress, significant alteration of DNA methylation was observed in waterlogging and drought conditions. Importantly, drought stress increased the global methylation level through an active *de novo* methylation while under waterlogging stress, the opposite scenario was observed. Drought stress has been investigated in various plants species at the epigenetic level. Our report is in agreement with the study of Labra et al. (2002) who showed a hypermethylation in pea root tip under water deficit. In contrast to our results, drought has been reported to decrease the level of total DNA methylation by 12.1% and 10.28% in rice and ryegrass, respectively (Wang et al., 2011; Tang et al., 2014). Similarly, a quantification of the genome wide cytosine methylation polymorphism using the MSAP analysis showed a predominant hypomethylation in a tolerant rice accession under drought stress (Gayacharan and Joel, 2013). Furthermore, Abid et al. (2017) recently noticed that drought stress reduces the methylation level in two faba bean genotypes, irrespective to their tolerance level. These contrasting reports imply that different methylation mechanisms for drought tolerance exist in plant species. Similar conclusions were drawn from the epigenetic responses of clover and hemp which decreased the methylation level under chromium stress but an active *de novo* synthesis of methylated cytosine was found to positively correlate with the intensity of the stress in the rape genome (Aina et al., 2004; Labra et al., 2004). Nonetheless, we deduced that hypermethylation under drought stress is a tolerance strategy in sesame and it would be valuable to compare in a future experiment the epigenetic alterations of (1) several sesame genotypes with contrasting levels of drought tolerance to understand the intra-species variation (Abid et al., 2017), (2) various plant species exhibiting contrasting responses to drought to uncover the inter-species variation (Aina et al., 2004).

Various abiotic stress have been investigated using MSAP markers including cold, drought, salt, aluminum stresses, however, waterlogging stress has been scarcely studied. Here, we showed that waterlogging stress in contrast to drought decreased the global methylation level in sesame. As a typical waterlogging-susceptible and drought-tolerant crop, we were expecting a contrasting epigenetic response of sesame to these abiotic stresses, hypothesis which has been confirmed in the present study. Is it possible that mimicking the effective drought epigenetic response by imposing a strong directed *de novo* DNA methylation during waterlogging stress will lead to stress tolerance? Unfortunately the mechanism underpinning the contrasting DNA methylation alteration under drought and waterlogging in sesame remains unclear. We suspected that DNA methyltransferases and DNA glucolsylase demethylase genes may be the master players in this mechanism (Gehring and Henikoff, 2007). In this case, the identification and manipulation of the key enzymes that control the level of methylation may assist in efforts to develop stress-tolerant sesame plants.

In nature, environmental stresses are rarely permanent and the ability of plants to fully recover after stress relief is an important component of stress resistance mechanisms (Peleg et al., 2011). Unfortunately, many studies overlooked the methylation patterns during the recovery phase. In this study, our data demonstrated a difference in the epigenetic behavior as a result of the recovery from drought and waterlogging damages. We showed that after recovery from drought, a reversion of the methylated sites took place and tended to reach the level under a stress-free condition. Similar to our observations, it was found that about 70% of the sites which exhibited drought-induced epigenetic methylation were demethylated when rice plants were allowed to recover (Wang et al., 2011). Hence, sesame plants were able to promptly recover an optimal physiological status after stress relief. However, during the waterlogging recovery phase, both demethylation and *de novo* methylation concomitantly occur in the sesame root. We inferred from that observation that waterlogging stress causes extensive damages which require prolonged physiological and morpho-anatomical adjustments to recover as compared to the drought recovery. Altogether, our MSAP analysis indicated that sesame has a divergent epigenetic program to respond to drought and waterlogging, which may explain its contrasting response to these major abiotic stresses.

DNA methylation and demethylation have been proven to be associated with gene expression leading to adaptive physiological and morphological responses to abiotic stress (Choi and Sano, 2007; Vining et al., 2012; Garg et al., 2015). Here, our comparative analysis of MSAP profiles and transcriptome data revealed a strong correlation between DNA methylation pattern and the regulation of the responsive-genes to drought and waterlogging in sesame. In particular, we deduced that *de novo* DNA methylation participates in the down-regulation of the gene expression while DNA demethylation results in the up-regulation of the gene expression. In addition, 90% of the sequenced polymorphic MSAP fragments corresponded to significantly and differentially expressed genes (DEGs) between treatments. Also, they were predominantly located in the promoter region of the protein-coding genes, which implies that DNA methylation and demethylation lead to the activation and inactivation of the transcriptional processes for specific genes related to drought and waterlogging response in sesame. In accordance with our results, Meng et al. (2016) by combining transcriptome and DNA methylation data from *Arabidopsis thaliana* concluded to a significant epigenetic contribution to gene expression regulation. Moreover, similar conclusions were drawn by Vining et al. (2012) who reported a negative correlation between methylation and transcription of gene at the genome wide level in *Populus trichocarpa*. Another interesting finding in this study concerns the quasi contrasting regulation of the shared DEGs between drought and waterlogging under stress and recovery stages. This important observation further shed light on the opposite molecular responses under drought and waterlogging treatments, directed by divergent epigenetic factors such as DNA methylation.

## 5. Conclusion

For the first time, the epigenetic responses of sesame to drought and waterlogging were revealed using the MSAP approach. Drought stress increased the methylation level while waterlogging induced a high demethylation, indicating an opposite epigenetic responsive program. At the recovery stage, drought-stressed plants promptly readjusted the methylation level through a strong demethylation. However, waterlogging stressed-plants required a coordinated methylation and demethylation activity during the recovery. A high degree of correlation was also observed between DNA methylation pattern and gene regulation with a high methylation associated with gene down-regulation while a strong demethylation was correlated with the up-regulation of the gene transcriptional activity. In summary, drought and waterlogging stress induce divergent DNA methylation patterns which influence gene regulation and lead to a contrasting response of sesame to these two abiotic stresses. Further investigations are required to better understand the key modulators of the DNA methylation in the sesame genome so as to modify epigenetic cascades and improve sesame productivity and yield under abiotic stress.

## Author’s contributions statement

Conception and design: KD, NC, DD, XZ; Production of the data: KD, MAM, QZ, MY, WL, RZ; Analysis and interpretation of the data: KD, LW, MAM; Drafting of the article: KD, MAM; Final approval of the article: KD, MAM, RZ, QZ, MY, NC, DD, LW, XZ.

## Ethical standards

The authors declare that the experiments comply with the current laws of the countries in which the experiments were performed.

## Acknowledgements

This work was funded by the China Agriculture research System (CARS-14) and the Agricultural Science and Technology Innovation Project of Agricultural Sciences (CAASASTIP-201-OCRI). The first two authors are grateful to the *Deutscher Akademischer Austausch Dienst* scholarship.

## Supplementary Tables

**Table S1.** List of the primers used for MSAP analysis

**Table S2.** The arrangement of all possible events explaining the sixteen four-bit binary MSAP codes (adapted from Bednarek et al., 2017)

**Table S3.** List of qRT-PCR primers

## References

Abid, G.D., Mingeot, Y., Muhovski, G., Mergeai, M., Aouida, S., Abdelkarim, I., Aroua, M., El Ayed, M., M’hamdi, K., Sassi, M., Jebara., 2017. Analysis of DNA methylation patterns associated with drought stress response in faba bean (*Vicia faba L.)* using methylation-sensitive amplification polymorphism (MSAP). Env. Exp. Bot. 142, 34–44.

Aina, R., Sgorbati, S., Santagostino, A., Labra, M., Ghiani, A., Citterio S., 2004. Specific hypomethylation of DNA is induced by heavy metals in white clover and industrial hemp. Physiol. Plant. 121, 472–80.

Al-Lawati, A., Al-Bahry, S., Victor, R., Al-Lawati, A.H., Yaish, M.W., 2016. Salt stress alters DNA methylation levels in alfalfa (*Medicago spp*). Genet. Mol. Res. 15, 1–16.

Ashikawa, I., 2001. Surveying CpG methylation at 5’-CCGG in the genomes of rice cultivars. Plant Mol. Biol. 45, 31–39.

Banerjee, A., Roychoudhury, A., 2017. Epigenetic regulation during salinity and drought stress in plants: Histone modifications and DNA methylation. Plant Gene 11, 199–204.

Baulcombe, D.C., Dean, C., 2014. Epigenetic regulation in plant responses to the environment. Cold Spring Harb. Perspect. Biol. 6, a019471.

Bedigian, D., 2004. History and lore of sesame in Southwest Asia. Econ. Bot. 58, 329–353.

Bednarek, P.T., Orłowska, R., Niedziela, A., 2017. A relative quantitative methylation-sensitive amplified polymorphism (MSAP) method for the analysis of abiotic stress. BMC Plant Biol. 17, 79.

Benbouza, H., Jacquemin, J.M., Baudoin, J.P., Mergeai, G., 2006. Optimization of a re- liable, fast, cheap and sensitive silver staining method to detect SSR markers in polyacrylamide gels. Biotechnol. Agron. Soc. Environ. 10, 77–81.

Boureima, S., Oukarroum, A., Diouf, M., Cissé, N., Van Damme, P., 2012. Screening for drought tolerance in mutant germplasm of sesame (*Sesamum indicum*) probing by chlorophyll a fluorescence. Env. Exp. Bot. 81, 37–43.

Chwialkowska, K., Korotko, U., Kosinska, J., Szarejko, I., Kwasniewski, M., 2017. Methylation sensitive amplification polymorphism sequencing (MSAP-Seq)- a method for high-throughput analysis of differentially methylated CCGG sites in plants with large genomes. Front. Plant Sci. 8, 2056.

Chwialkowska, K., Nowakowska, U., Mroziewicz, A., Szarejko, I., Kwasniewski, M., 2016. Water-deficiency conditions differently modulate the methylome of roots and leaves in barley (*Hordeum vulgare L*.). J. Exp. Bot. 67, 1109–1121.

Cokus, S.J., Feng, S.H., Zhang, X., Chen, Z., Merriman, B., Haudenschild, C.D, Pradhan, S., Nelson, S.F., Pellegrini, M., Jacobsen, S.E., 2008. Shotgun bisulfite sequencing.g of the *Arabidopsis* genome reveals DNA methylation patterning. Nature 452, 215–219

Dai, L.F., Chen, Y.L., Luo, X.D., Wen, X.F., Cui, F.L., Zhang, F.T., Zhou, Y., Xie, J.K., 2015. Level and pattern of DNA methylation changes in rice cold tolerance introgression lines derived from *Oryza rufipogon* Griff. Euphytica 205, 73–83.

Dossa, K., Diouf, D., Wang, L., Wei, X., Zhang, Y., Niang, M., Fonceka, D., Yu, J., Mmadi, M.A, Yehouessi, LW., Liao, B., Zhang, X., Cisse, N., 2017a. The emerging oilseed crop *Sesamum indicum* enters the “Omics” era. Front. Plant Sci. 8, 1154.

Dossa, K., Li, D., Wang, L., Zheng, X., Liu, A., Yu, J., Wei, X., Zhou, R., Fonceka, D., Diouf, D., et al., 2017d. Transcriptomic, biochemical and physio-anatomical investigations shed more light on responses to drought stress in two contrasting sesame genotypes. Sci. Rep. 7, 8755.

Dossa, K., Li, D., Wang, L., Zheng, X., Yu, J., Wei, X., et al., 2017c. Dynamic transcriptome landscape of sesame (*Sesamum indicum L*.) under progressive drought and after rewatering. Genom. Data 11, 122–124.

Dossa, K., Wei, X., Zhang, Y., Fonceka, D., Yang, W., Diouf, D., et al., 2016. Analysis of genetic diversity and population structure of sesame accessions from Africa and Asia as major centers of its cultivation. Genes 7, 14.

Dossa, K., Yehouessi, W.L., Likeng, B. C., Diouf, D., Liao, B., Zhang, X., Cissé N., Bell J. M., 2017b. Comprehensive screening of West and Central African sesame accessions for drought tolerance probing by agro-morphological, physiological, biochemical and nutritional traits. Agronomy 7, 83.

Dowen, R.H., Pelizzola, M., Schmitz, R.J., Lister, R., Dowen, J.M., Nery, J.R., et al., 2012. Widespread dynamic DNA methylation in response to biotic stress. Proc. Natl. Acad. Sci. U.S.A. 109, 12858–12859.

Farooq, M., Wahid, A., Kobayashi, N., Fujita, D., Basra, S.M.A., 2009. Plant drought stress: effects, mechanisms and management. Agron. Sustain. Dev. 29, 185–212.

Fulneček, J., Kovařík, A., 2014. How to interpret Methylation Sensitive Amplified Polymorphism (MSAP) profiles? BMC Genet. 15, 2.

Garg, R., Narayana Chevala, V.V.S., Shankar, R., Jainb, M., 2015. Divergent DNA methylation patterns associated with gene expression in rice cultivars with contrasting drought and salinity stress response. Sci. Rep. 5, e14922.

Gayacharan, A., Joel J., 2013. Epigenetic responses to drought stress in rice (*Oryza sativa L*.). Physiol. Mol. Biol. Plants 19, 379–387.

Gehring, M., Henikoff, S., 2007. DNA methylation dynamics in plant genomes. Biochim. Biophys. Acta 1769, 276e286.

Goll, M.G., Kirpekar, F., Maggert, K.A., Yoder, J.A., Hsieh, C.L., Zhang, X., Golic, K.G., Jacobsen, S.E., Bestor, T.H., 2006. Methylation of tRNAAsp by the DNA methyltransferase homolog Dnmt2. Science 311, 395–8.

Hassanzadeh, M., Ebadi, M., Panahyan-e-Kivi, M., Jamaati-e-Somarin, S.H., Saeidi, M., Zabihi-e-Mahmoodabad, R., 2009. Evaluation of drought stress on relative water content and chlorophyll content of sesame (*Sesamum indicum L*.) genotypes at early flowering stage. Res. J. Environ. Sci. 3, 345–350.

Karan, R., DeLeon, T., Biradar, H., Subudhi, P.K., 2012. Salt stress induced variation in DNA methylation pattern and its influence on gene expression in contrasting rice genotypes. PLoS One 7, e40203.

Kim, D., Langmead, B., Salzberg, S.L., 2015. HISAT: a fast spliced aligner with low memory requirements. Nat. Method. 12, 357–360.

Labra, M., Ghiani, A., Citterio, S., Sgorbati, S., et al., 2002. Analysis of cytosine methylation pattern in response to water deficit in pea root tips. Plant Biol. 4, 694–699.

Labra, M., Grassi, F., Imazio, S., et al., 2004. Genetic and DNA-methylation changes induced by potassium dichromate in *Brassica napus L*. Chemosphere 54, 1049–58.

Lalitha, S., 2000. Primer premier 5. Biotechnol. Softw. Internet Rep. 1, 270–272.

Langham, D.R., 2007. “Phenology of sesame,” In: J. Janick and A. Whipkey (ed.), Issues in New Crops and New Uses, ASHS Press (pp. 144–182), Alexandria, VA.

Livak, K.J., Schmittgen, T.D., 2001. Analysis of relative gene expression data using real-time quantitative PCR and the 2(-Delta Delta C (T)) Method. Methods 25, 402–408.

Marconi, G., Pace, R., Traini, A., Raggi, L., Lutts, S., et al., 2013. Use of MSAP markers to analyse the effects of salt stress on DNA methylation in rapeseed (*Brassica napus var. oleifera*). PLoS ONE 8, e75597.

Matthes, M., Singh, R., Cheah, S.C., Karp, A., 2001. Variation in oil palm (*Elaeis guineensis Jacq.*) tissue culture-derived regenerants revealed by AFLPs with methylation-sensitive enzymes. Theor. Appl. Genet. 102, 971–979.

Meng, D., Dubin., M., Zhang, P., Osborne, E.J., Stegle, O., Clark, R.M., Nordborg, M., 2016. Limited contribution of DNA methylation variation to expression regulation in *Arabidopsis thaliana*. PLoS Genet. 12, e1006141.

Pan, L., Liu, X., Wang, Z., 2012. Comparative DNA methylation analysis of powdery mildew susceptible and resistant near-isogenic lines in common wheat. Life Sci. J. 10, 2073–2083.

Pathak, N., Rai, A. K., Kumari, R., Bhat, K.V., 2014. Value addition in sesame: a perspective on bioactive components for enhancing utility and profitability. Pharmacogn. Rev. 8, 147–155.

Paun, O., Bateman, R.M., Fay, M.F., Hedrén, M., Civeyrel, L., Chase, M., 2010. Stable epigenetic effects impact adaptation in allopolyploid orchids (*Dactylorhiza: Orchidaceae*). Mol. Biol. Evol. 27, 2465–2473.

Peleg, Z., Apse, M.P., Blumwald, E., 2011. Engineering salinity and water-stress tolerance in crop plants: getting closer to the field. In: I. Turkan. Adv. Bot. Res. Academic Press. 57, 405–443.

Pertea, M., Kim, D., Pertea, G., Leek, J.T., Salzberg, S.L., 2016. Transcript-level expression analysis of RNA-seq experiments with HISAT, StringTie and Ballgown. Nat. Protoc. 11, 1650–1667.

Portis, E., Acquadro, A., Comino, C., Lanteri, S., 2004. Analysis of DNA methylation during germination of pepper (*Capsicum annuum L*.) seeds using methylation-sensitive amplification polymorphism (MSAP). Plant Sci. 166, 169–178.

Reyna-López, G.E., Simpson, J., Ruiz-Herrera, J., 1997. Differences in DNA methylation patterns are detectable during the dimorphic transition of fungi by amplification of restriction polymorphisms. Mol. Gen. Genet. 253, 703–710.

Ruiz-Garcia, L., Cervera, M.T., Martinez-Zapater, J.M., 2005. DNA methylation increases throughout *Arabidopsis* development. Planta 222, 301–306.

Saeed, A.I., Bhagabati, N.K., Braisted, J.C., Liang, W., Sharov, V., Howe, E.A., Li, J.W., Thiagarajan, M., White, J.A., Quackenbush, J., 2006. TM4 microarray software suite. Method Enzymol. 411, 134–193.

Schulz, B., Eckstein, R.L., Durka, W., 2013. Scoring and analysis of methylation-sensitive amplification polymorphisms for epigenetic population studies. Mol. Ecol. Resour. 13, 642–53.

Sha, A.H., Lin, X.H., Huang, J.B., Zhang, D.P., 2005. Analysis of DNA methylation related to rice adult plant resistance to bacterial blight based on methylation-sensitive AFLP (MSAP) analysis. Mol. Genet. Genom. 273, 484–490.

Shan, X., Wang, X., Yang, G., Wu, Y., Su, S., Li, S., Li, H., Yuan, Y., 2013. Analysis of the site-specific DNA methylation and its association with drought tolerance in response to cold stress based on methylation-sensitive amplified polymorphisms. J. Plant Biol. 56, 32–38.

Sun, J., Rao, Y., Le, M., Yan, T., Yan, X., Zhou, H., 2010. Effects of drought stress on sesame growth and yield characteristics and comprehensive evaluation of drought tolerance. Chin. J. Oil Crop Sci. 32, 525–533.

Tang, X.M., Tao, X., Wang, Y., Ma, D.W., Li, D., Yang, H., Ma, X.R., 2014. Analysis of DNA methylation of perennial under drought using the methylation-sensitive amplification polymorphism (MSAP) technique. Mol. Genet. Genom. 289, 1075–1084.

Tarazona, S., Garcia-Alcalde, F., Dopazo, J., Ferrer, A., Conesa, A., 2011. Differential expression in RNA-seq: a matter of depth. Gen. Res. 21, 2213–2223.

Trapnell C., Williams B.A., Pertea G., Mortazavi A., Kwan G., van Baren M.J., et al., 2010. Transcript assembly and quantification by RNA-Seq reveals unannotated transcripts and isoform switching during cell differentiation. Nat. Biotechnol. 28, 511–5.

Vining, K. J., Pomraning, K.R., Wilhelm, L.J., Priest, H.D., Pellegrini, M., Mockler, T.C., Freitag, M. Strauss, S.H., 2012. Dynamic DNA cytosine methylation in the *Populus trichocarpa* genome: tissue-level variation and relationship to gene expression. BMC Genom. 13, 27.

Wang, L., Li, D., Zhang, Y., Gao, Y., Yu, J., Wei, X., et al., 2016a. Tolerant and susceptible sesame genotypes reveal waterlogging stress response patterns. PLoS ONE 11, e0149912.

Wang, L., Xia, Q., Zhang, Y., Zhu, X., Zhu, X., Li, D., et al., 2016b. Updated sesame genome assembly and fine mapping of plant height and seed coat color QTLs using a new high-density genetic map. BMC Genom. 17, 31.

Wang, L., Yu, J., Li, D., Zhang, X., 2014. Sinbase: an integrated database to study genomics, genetics and comparative genomics in *Sesamum indicum*. Plant Cell Physiol. 56, e2.

Wang, L., Zhang, Y., Qi, X., Li, D., Wei, W., Zhang, X., 2012. Global gene expression responses to waterlogging in roots of sesame (*Sesamum indicum L.*). Acta Physiol. Plant. 34, 2241–2249.

Wang, W., Huang, F., Qin, Q., Zhao, X., Li, Z., Fu. B., 2015. Comparative analysis of DNA methylation changes in two rice genotypes under salt stress and subsequent recovery. Biochem. Biophys. Res. Commun. 465, 790e796.

Wang, W.S., Pan, Y.J., Zhao, X.Q., Dwivedi, D., Zhu, L.H., et al., 2011. Drought induced site-specific DNA methylation and its association with drought tolerance in rice (*Oryza sativa L*.). J. Exp. Bot. 62, 1951–1960.

Xia, H., Huang, W.X., Xiong, J., Tao, T., Zheng, X. G., Wei, H.B., et al., 2016. Adaptive epigenetic differentiation between upland and lowland rice ecotypes revealed by methylation-sensitive amplified polymorphism. PLoS ONE 11, e0157810.

Xiong, L.Z., Xu, C.G., Saghai Maroof, M.A., Zhang, Q., 1999. Patterns of cytosine methylation in an elite rice hybrid and its parental lines, detected by a methylation-sensitive amplification polymorphism technique. Mol. Gen. Genet. 261, 439–46.

You, J., Wang, Y., Zhang, Y., Dossa, K., Li, D., Zhou, R., Wang, L., Zhang, X., 2018. Genome-wide identification and expression analyses of genes involved in raffinose accumulation in sesame. Sci. Rep. 8, 4331.

Zhang, P.Y, Wang, J.G, Geng, Y.P, Dai, J.R, Zhong, Y., Chen, Z.Z, Zhu, K., Wang, X.Z., Chen, S.Y., 2015. MSAP-based analysis of DNA methylation diversity in tobacco exposed to different environments and at different development phases. Biochem. Syst. Ecol. 62, 249–260.

Zheng X., Pontes, O., Zhu, J., Miki, D., Zhang, F., Li, W.X., Iida, K., Kapoor, A., Pikaard, C.S., Zhu, J.K., 2008. ROS3 is an RNA-binding protein required for DNA demethylation in *Arabidopsis*. Nature 30, 259–62.

